# A mini-atlas of gene expression for the domestic goat (*Capra hircus*) reveals transcriptional differences in immune signatures between sheep and goats

**DOI:** 10.1101/711127

**Authors:** Charity Muriuki, Stephen J. Bush, Mazdak Salavati, Mary E.B. McCulloch, Zofia M. Lisowski, Morris Agaba, Appolinaire Djikeng, David A. Hume, Emily L. Clark

## Abstract

Goats (*Capra hircus*) are an economically important livestock species providing meat and milk across the globe. They are of particular importance in tropical agri-systems contributing to sustainable agriculture, alleviation of poverty, social cohesion and utilisation of marginal grazing. There are excellent genetic and genomic resources available for goats, including a highly contiguous reference genome (ARS1). However, gene expression information is limited in comparison to other ruminants. To support functional annotation of the genome and comparative transcriptomics we created a mini-atlas of gene expression for the domestic goat. RNA-Seq analysis of 22 transcriptionally rich tissues and cell-types detected the majority (90%) of predicted protein-coding transcripts and assigned informative gene names to more than 1000 previously unannotated protein-coding genes in the current reference genome for goat (ARS1). Using network-based cluster analysis we grouped genes according to their expression patterns and assigned those groups of co-expressed genes to specific cell populations or pathways. We describe clusters of genes expressed in the gastro-intestinal tract and provide the expression profiles across tissues of a subset of genes associated with functional traits. Comparative analysis of the goat atlas with the larger sheep gene expression atlas dataset revealed transcriptional differences between the two species in macrophage-associated signatures. The goat transcriptomic resource complements the large gene expression dataset we have generated for sheep and contributes to the available genomic resources for interpretation of the relationship between genotype and phenotype in small ruminants.

## Introduction

Goats (*Capra hircus*) are an important source of meat and milk globally. They are an essential part of sustainable agriculture in low and middle-income countries, representing a key route out of poverty particularly for women. Genomics-enabled breeding programmes for goats are currently implemented in the UK and France with breeding objectives including functional traits such as reproductive performance and disease resistance (Larroque et al., 2016; Pulina et al., 2018). The International Goat Genomics Consortium (IGGC) (http://www.goatgenome.org) has provided extensive genetic tools and resources for goats including a 52K SNP chip (Tosser-Klopp et al., 2014), a functional SNP panel for parentage assessment and breed assignment (Talenti et al., 2018) and large-scale genotyping datasets characterising global genetic diversity (Stella et al., 2018). In 2017 a highly contiguous reference genome for goat (ARS1) was released (Bickhart et al., 2017; Worley, 2017). Advances in genome sequencing technology, particularly the development of long-read and single-molecule sequencing, meant that the ARS1 assembly was a considerable improvement in quality and contiguity from the previous whole genome shotgun assembly (CHIR_2.0) (Dong et al., 2013). In 2018 the ARS1 assembly was released on the Ensembl genome portal (Zerbino et al., 2018) (https://www.ensembl.org/Capra_hircus/Info/Index) greatly facilitating the utility of the new assembly and providing a robust set of gene models for goat.

RNA-Sequencing (RNA-Seq) has transformed the analysis of gene expression from the single-gene to the whole genome allowing visualisation of the entire transcriptome and defining how we view the transcriptional control of complex traits in livestock (reviewed in (Wickramasinghe et al., 2014)). Using RNA-Seq we generated a large-scale high-resolution atlas of gene expression for sheep (Clark et al., 2017). This dataset included RNA-Seq libraries from all organ systems and multiple developmental stages, providing a model transcriptome for ruminants. Analysis of the sheep gene expression atlas dataset indicated we could capture approximately 85% of the transcriptome by sampling twenty ‘core’ tissues and cell types (Clark et al., 2017). Given the close relationship between sheep and goats, there seemed little purpose in replicating a resource on the same scale. Our aim with the goat mini-atlas project, which we present here, was to produce a smaller, cost-effective, atlas of gene expression for the domestic goat based on transcriptionally rich tissues from all the major organ systems.

In the goat genome there are still many predicted protein-coding and non-coding genes for which the gene model is either incorrect or incomplete, or where there is no informative functional annotation. For example, in the current goat reference genome, ARS1 (Ensembl release 97), 33% of the protein-coding genes are identified only with an Ensembl placeholder ID. Many of these unannotated genes are likely to have important functions. Using RNA-Seq data we can annotate them and assign function (Krupp et al., 2012). With datasets of a sufficient size, genes form co-expression clusters, which can either be ubiquitous, associated with a cellular process or be cell-/tissue specific. This information can then be used to associate a function with genes co-expressed in the same cluster, a method of functional annotation known as the ‘guilt by association principle’ (Oliver, 2000). Using this principle with the sheep gene expression atlas dataset we were able to annotate thousands of previously unannotated transcripts in the sheep genome (Clark et al., 2017). By applying this rationale to the goat mini-atlas dataset we were able to do the same for the goat genome.

The goat mini-atlas dataset that we present here was used by Ensembl to create the initial gene build for ARS1 (Ensembl release 92). A high-quality functional annotation of existing reference genomes can help considerably in our understanding of the transcriptional control of functional traits to improve the genetic and genomic resources available, inform genomics enabled breeding programmes and contribute to further improvements in productivity. The entire dataset is available in a number of formats to support the livestock genomics research community and represents an important contribution to the Functional Annotation of Animal Genomes (FAANG) project (Andersson et al., 2015; FAANG, 2017; Harrison et al., 2018).

This study is the first global analysis of gene expression in goats. Using the goat mini-atlas dataset we describe large clusters of genes associated with the gastrointestinal tract and macrophages. Species specific differences in response to disease, or other traits, are likely to be reflected in gene expression profiles. Sheep and goats are both small ruminant mammals and are similar in their physiology. They also share susceptibility to a wide range of viral, bacterial, parasitic and prion pathogens, including multiple potential zoonoses (Sherman, 2011), but there have been few comparisons of relative susceptibility or pathology between the species to the same pathogen, nor the nature of innate immunity. To reveal transcriptional similarities and differences between sheep and goats we have performed a comparative analysis of species-specific gene expression by comparing the goat mini-atlas dataset with a comparable subset of data from the sheep gene expression atlas (Clark et al., 2017). We also use the goat mini-atlas dataset to examine the expression of candidate genes associated with functional traits in goats and link these with allele-specific expression (ASE) profiles across tissues, using a robust methodology for ASE profiling (Salavati et al., 2019). The goat mini-atlas dataset and the analysis we present here provide a foundation for identifying the regulatory and expressed elements of the genome that are driving functional traits in goats.

## Methods

### Animals

Tissue and cell samples were collected from six male and one female neonatal crossbred dairy goats at six days old. The goats were sourced from one farm and samples were collected at a local abattoir within 1 hour of euthanasia.

### Tissue Collection

The tissue samples were excised post-mortem within one hour of death, cut into 0.5cm diameter segments and transferred into RNAlater (Thermo Fisher Scientific, Waltham, USA) and stored at 4°C for short-term storage. Within one week, the tissue samples were removed from the RNAlater, transferred to 1.5ml screw cap cryovials and stored at −80°C until RNA isolation. Alveolar macrophages (AMs) were isolated from two male goats by broncho-alveolar lavage of the excised lungs using the method described for sheep in (Clark et al., 2017), except using 20% heat-inactivated goat serum (G6767, Sigma Aldrich), and stored in TRIzol (15596018; Thermo Fisher Scientific) for RNA extraction. Similarly bone marrow derived macrophages (BMDMs) were isolated from 10 ribs from 3 male goats and frozen down for subsequent stimulation with lipopolysaccharide (LPS) (*Salmonella enterica* serotype minnesota Re 595 (L9764; Sigma-Aldrich)) using the method described in (Clark et al., 2017; Young et al., 2018) with homologous serum. Details of all the samples collected are included in Table 1.

**Table 1:** Details of samples included in the goat mini-atlas.

| **Tissue/Cell type** | **Organ System** | **No. of replicates** | **Sex** |
| --- | --- | --- | --- |
| **Adrenal gland** | Endocrine | 4 | male |
| **Alveolar macrophage** | Immune | 2 | male |
| **BMDM - LPS (0 hours)** | Immune | 3 | male |
| **BMDM + LPS (7 hours)** | Immune | 3 | male |
| **Cerebellum** | Nervous system | 2 | male |
| **Colon large** | GI tract | 4 | male |
| **Fallopian tube** | Reproductive system (female) | 1 | female |
| **Frontal lobe cortex** | Nervous system | 2 | male |
| **Ileum and Peyer's patches** | GI tract | 2 | male |
| **Kidney cortex** | Endocrine | 4 | male |
| **Liver** | Endocrine | 4 | male |
| **Ovary** | Reproductive system (female) | 1 | female |
| **Rumen** | Gastrointestinal tract | 2 | male |
| **Skeletal muscle - longissimus dorsi** | Musculo-skeletal | 3 | male |
| **Skin** | Integumentary | 4 | male |
| **Spleen** | Immune | 3 | male |
| **Testes** | Reproductive system (male) | 4 | male |
| **Thymus** | Immune | 4 | male |
| **Uterine horn** | Reproductive system (female) | 1 | female |
| **Uterus** | Reproductive system (female) | 1 | female |

### RNA extraction

RNA was extracted from tissues and cells as described in (Clark et al., 2017). For each RNA extraction from tissues approximately 60mg of tissue was processed. Tissue samples were first homogenised in 1ml of TRIzol (15596018; Thermo Fisher Scientific) with CK14 (432–3751; VWR, Radnor, USA) tissue homogenising ceramic beads on a Precellys Tissue Homogeniser (Bertin Instruments; Montigny-le-Bretonneux, France) at 5000 rpm for 20 sec. Cell samples which had previously been collected in TRIzol (15596018; Thermo Fisher Scientific) were mixed by pipetting to homogenise. Homogenised (cell/tissue) samples were then incubated at room temperature for 5 min to allow complete dissociation of the nucleoprotein complex, 200µl BCP (1-bromo-3-chloropropane) (B9673; Sigma Aldrich) was added, then the sample was shaken vigorously for 15 sec and incubated at room temperature for 3 min. The sample was centrifuged for 15 min at 12,000 x *g*, at 4°C for 3 mins to separate the homogenate into a clear upper aqueous layer. The homogenate was then column purified to remove DNA and trace phenol using a RNeasy Mini Kit (74106; Qiagen Hilden, Germany) following the manufacturer’s instructions (RNeasy Mini Kit Protocol: Purification of Total RNA from Animal Tissues, from step 5 onwards). An on-column DNase treatment was performed using the Qiagen RNase-Free DNase Set (79254; Qiagen Hilden, Germany). The sample was eluted in 30ul of RNase free water and stored at −80°C prior to QC and library preparation. RNA integrity (RIN^e^) was estimated on an Agilent 2200 TapeStation System (Agilent Genomics, Santa Clara, USA) using the RNA Screentape (5067–5576; Agilent Genomics) to ensure RNA quality was of RIN^e^ > 7. RIN^e^ and other quality control metrics for the RNA samples are included in Supplementary Table S1.

### RNA-Sequencing

RNA-Seq libraries were prepared by Edinburgh Genomics (Edinburgh Genomics, Edinburgh, UK) and run on the Illumina HiSeq 4000 sequencing platform (Illumina, San Diego, USA). Strand-specific paired-end reads with a fragment length of 75bp were generated for each sample using the standard Illumina TruSeq mRNA library preparation protocol (poly-A selected) (Ilumina; Part: 15031047 Revision E). Libraries were sequenced at a depth of either >30 million reads per sample for the tissues and AMs, or >50 million reads per sample for the BMDMs.

### Data Processing

The RNA-Seq data processing methodology and pipelines are described in detail in (Clark et al., 2017). Briefly, for each tissue a set of expression estimates, as transcripts per million (TPM), were obtained using the alignment-free (technically, ‘pseudo-aligning’) transcript quantification tool Kallisto (Bray et al., 2016), the accuracy of which depends on a high quality index (reference transcriptome). In order to ensure an accurate set of gene expression estimates we used a ‘two-pass’ approach to generate this index.

We first ran Kallisto on all samples using as its index the ARS1 reference transcriptome available from Ensembl (ftp://ftp.ensembl.org/pub/release-95/fasta/capra_hircus/cdna/Capra_hircus.ARS1.cdna.all.fa.gz). We then parsed the resulting data to revise this index. This was for two reasons: i) in order to include, in the second index, those transcripts that should have been present but were missing (i.e. where the reference annotation was incomplete), and ii) to remove those transcripts that were present but should not have been (i.e. where the reference annotation was poor quality and a spurious model had been introduced). For i) we obtained the subset of reads that Kallisto could not (pseudo)align, assembled those *de novo* into putative transcripts, then retained each transcript only if it could be robustly annotated (by, for instance, encoding a protein similar to one of known function) and showed coding potential. For ii), we identified those transcripts in the reference transcriptome for which no evidence of expression could be found in any of the samples from the goat mini-atlas. These were then discarded from the index and the revised index was used for a second ‘pass’ with Kallisto, generating higher-confidence expression level estimates.

We complemented the Kallisto alignment-free method with a reference-guided alignment-based approach to RNA-Seq processing, using the HISAT aligner (Kim et al., 2015) and StringTie assembler (Pertea et al., 2015). This approach was highly accurate when mapping to the (ARS1) annotation on NCBI (ftp://ftp.ncbi.nlm.nih.gov/genomes/all/GCF/001/704/415/GCF_001704415.1_ARS1/GCF_001704415.1_ARS1_rna.fna.gz), precisely reconstructing almost all exon (96%) and transcript (76%) models (Supplementary Table S2). We used the HISAT/StringTie output to validate the set of transcripts used to generate the Kallisto index. Unlike alignment-free methods, HISAT/StringTie can be used to identify novel transcript models, particularly for ncRNAs, which we have described separately in (Bush et al., 2018b). Details of all novel transcript models detected are included in Supplementary Table S3.

### Data Validation

To identify any spurious samples which could have been generated during sample collection, RNA extraction or library preparation, we generated a sample-to-sample correlation of the gene expression estimates from Kallisto, in Graphia Professional (Kajeka Ltd, Edinburgh, UK).

### Network cluster analysis

Network cluster analysis of the goat gene mini-atlas dataset was performed using Graphia Professional (Kajeka Ltd, Edinburgh, UK) (Livigni et al., 2018). In brief, similarities between individual gene expression profiles were determined by calculating a Pearson correlation matrix for both gene-to-gene and sample-to-sample comparisons, and filtering to remove relationships where *r* < 0.83. A network graph was constructed by connecting the remaining nodes (transcripts) with edges (where the correlation exceeded the threshold value). The resultant graph was interpreted by applying the Markov Cluster algorithm (MCL) at an inflation value (which determines cluster granularity) of 2.2. The local structure of the graph was then examined visually. Transcripts with robust co-expression patterns, i.e. related functions, clustered together forming sets of tightly interlinked nodes. The principle of ‘guilt by association’ was then applied, to infer the function of unannotated genes from genes within the same cluster (Oliver, 2000). Expression profiles for each cluster were examined in detail to understand the significance of each cluster in the context of the biology of goat tissues and cells. Clusters 1 to 30 were assigned a functional ‘class’ and ‘sub-class’ manually by first determining if multiple genes within a cluster shared a similar biological function based on GO term enrichment using the Bioconductor package ‘topGO’ (Alexa and Rahnenfuhrer, 2010).

### Comparative analysis of gene expression in macrophages in sheep and goats

To compare transcriptional differences in the immune response between the two species we focused our analysis on the macrophage populations (AMs and BMDMs). For this analysis we used a subset of data from our sheep gene expression atlas for AMs and BMDMs (+/- LPS) from three male sheep (Clark et al., 2017) (Supplementary Dataset S1).

For AMs we compared the gene level expression estimates from the two male goats and three male sheep using edgeR v3.20.9 (Robinson et al., 2010). Only genes with the same gene name in both species, expressed at a raw read count of more than 10, FDR<10%, an FDR adjusted p-value of <0.05, and Log2FC of >=2, in both goat and sheep, were included in the analysis.

Differential expression analysis using edgeR (Robinson et al., 2010) was also performed for sheep and goat BMDMs (+/-) LPS separately, using the filtration criteria described above for AMs, to compile a list of genes for each species that were up or down regulated in response to LPS. These lists were then compared using the R package dplyr (Wickham et al., 2018) with system query language syntax. Each list was merged based on GENE_ID using the *inner_join* function to only return the observations that overlapped between goat and sheep (i.e. genes which had corresponding annotations in both species). A dissimilarity index (Dis_Index) was then calculated by taking the absolute difference of the Log2 fold change (Log2FC) between sheep and goat using the formula:

ABS(Log2FC Sheep-Log2FC Goat)

A high Dis_Index indicated that a gene was differently regulated in goat and sheep.

### Allele-specific expression

To measure allele-specific expression (ASE), across tissues and cell-types from the goat mini-atlas we used the method described in (Salavati et al., 2019). Briefly, BAM files from the RNA-Seq data, were mapped to the ARS1 top level DNA fasta track from Ensembl v96, using HISAT2 as described in (Clark et al., 2017). Any reference mapping bias was removed using WASP v0.3.1 (van de Geijn et al., 2015) and the resultant BAM files processed using the Genome Analysis Tool Kit (GATK) to produce individual VCF files. The ASEreadCounter tool in GATK v3.8 was used to obtain raw counts of the allelic expression profile in the dataset. These raw counts were then tested for imbalance (using a modified negative-beta bionomial test at gene level) at all heterozygote loci (i.e. ASE = Counts _RefAllele_/(Counts _RefAllele_+ Counts _AltAllele_) within the boundaries of the gene using the R package GeneiASE (Edsgärd et al., 2016).

## Results and Discussion

### Scope of the goat mini-atlas dataset, sequencing depth and coverage

The goat mini-atlas dataset includes 54 mRNA-Seq (poly-A selected) 75bp paired-end libraries. Details of the libraries generated including the age and sex of the animals, the tissues and cell types sampled, and the number of biological replicates per sample are summarised in Table 1. Gene level expression estimates, for the goat mini-atlas, are provided as unaveraged (Supplementary Dataset S2) and averaged across biological replicates (Supplementary Dataset S3) files.

Approximately 8.7×10^8^ paired end sequence reads were generated in total. Following data processing with Kallisto (Bray et al., 2016), a total of 18,528 unique protein coding genes had detectable expression (TPM>1), representing 90% of the reference transcriptome (Bickhart et al. 2017). From the set of 17 tissues and 3 cell types we sampled we were able to detect approximately 90% of protein coding genes providing proof of concept that the mini-atlas approach is useful for global analysis of transcription. The average percentage of transcripts detected per tissue or cell type was 66%, ranging from 54% in alveolar macrophages, which had the lowest to 72% in testes, which had the highest. The percentage of protein coding genes detected per each tissue is included in Table 2. Although we included uterine horn as well as uterus and both stimulated and unstimulated BMDMs, our analysis suggests that including only one tissue/cell of a similar type would be the most economical approach to generating a mini-atlas of gene expression for functional annotation.

**Table 2:** The percentage of protein coding genes detected per tissue in the goat mini-atlas dataset.

| **Tissue** | **Average no. of protein-coding genes expressed (TPM > 1) in this tissue** | **% of protein-coding genes expressed (TPM > 1) in this tissue** |
| --- | --- | --- |
| **Adrenal gland** | 14585 | 68.34 |
| **Alveolar macrophage** | 11533 | 54.04 |
| **BMDM - LPS (0 hours)** | 13253 | 62.1 |
| **BMDM + LPS (7 hours)** | 13042 | 61.11 |
| **Cerebellum** | 14959 | 70.09 |
| **Colon large** | 14736 | 69.04 |
| **Fallopian tube** | 14390 | 67.42 |
| **Frontal lobe cortex** | 14757 | 69.14 |
| **Ileum & Peyer's patches** | 15268 | 71.54 |
| **Kidney cortex** | 15223 | 71.33 |
| **Liver** | 13497 | 63.24 |
| **Ovary** | 14251 | 66.77 |
| **Rumen** | 13642 | 63.92 |
| **Skeletal muscle - longissimus dorsi** | 12276 | 57.52 |
| **Skin** | 14892 | 69.77 |
| **Spleen** | 14659 | 68.68 |
| **Testes** | 15359 | 71.96 |
| **Thymus** | 14484 | 67.86 |
| **Uterine horn** | 14298 | 66.99 |
| **Uterus** | 14298 | 66.99 |

Approximately 2,815 (13%) of the total 21,343 protein coding genes in the goat reference transcriptome had no detectable expression in the goat mini-atlas dataset. These transcripts are likely to be either tissue specific to tissues and cell-types that were not sampled here (including lung, heart, pancreas and various endocrine organs) rare or not detected at the depth of coverage used. The large majority of these transcripts were detected in the much larger sheep atlas, and their likely expression profile can be inferred from the sheep. In addition, for the goat mini-atlas unlike the sheep gene expression atlas we only included neonatal animals so transcripts that were highly developmental stage-specific in their expression pattern would also not be detected. A list of all undetected genes is included in Supplementary Table S4 and undetected transcripts in Supplementary Table S5.

### Gene Annotation

The proportion of transcripts per biotype (lncRNA, protein coding, pseudogene, etc), with detectable expression (TPM >1) in the goat mini-atlas relative to the ARS1 reference transcriptome, on Ensembl is summarised at the gene level in Supplementary Table S6 and at the transcript level in Supplementary Table S7. Of the 21,343 protein coding genes in the ARS1 reference transcriptome 7036 (33%) had no informative gene name. Whilst the Ensembl annotation will often identify homologues of a goat gene model, the automated annotation genebuild pipeline used to assign gene names and symbols is conservative. Using the annotation pipeline we described in (Clark et al., 2017) we were able to use the goat mini-atlas dataset to assign an informative gene name to 1114 previously un-annotated protein coding genes in ARS1. These genes were annotated by reference to the NCBI non-redundant (nr) peptide database v94 (Pruitt et al., 2007). A shortlist containing a conservative set of gene annotations to HGNC (HUGO Gene Nomenclature Committee) gene symbols, is included in Supplementary Table S8. Supplementary Table S9 contains the full list of genes annotated using the goat mini-atlas dataset and our annotation pipeline. Many unannotated genes can be associated with a gene description, but not necessarily an HGNC symbol; these are also listed in Supplementary Table S10. We manually validated the assigned gene names on the full list using network cluster analysis and the “guilt by association” principle.

### Network Cluster Analysis

Network cluster analysis of the goat gene expression atlas was performed using Graphia Professional (Kajeka Ltd, Edinburgh UK), a network visualisation tool (Livigni et al., 2018). The goat mini-atlas unaveraged TPM estimates (Supplementary Dataset S2) were used for network cluster analysis. We first generated a sample-to-sample graph (r=0.75, MCL=2.2) Supplementary Fig S1, which verified that the correlation between biological replicates was high and that none of the samples were spurious. We then generated a gene-to-gene network graph (Fig 1), with a Pearson correlation coefficient of r=0.83, that comprised 16,172 nodes (genes) connected by 1,574,259 edges. The choice of Pearson correlation threshold is optimised within the Graphia program to maximise the number of nodes (genes) included whilst minimising the number of edges. By applying the MCL (Markov Clustering) algorithm at an inflation value (which determines cluster granularity) of 2.2, the gene network graph separated into 75 distinct co-expression clusters, with the largest cluster (cluster 1) comprising of 1795 genes. Genes found in the top 30 largest clusters are listed in Supplementary Table S11. Clusters 1 to 20 (numbered in order of size, largest to smallest) were annotated manually and assigned a functional ‘class’ (Table 3). These functional classes were assigned based on GO term enrichment (Alexa and Rahnenfuhrer, 2010) for molecular function and biological process (Supplementary Table S12). Assignment of functional class was further validated by visual inspection of expression pattern and comparison with functional groupings of genes observed in the sheep gene expression atlas (Clark et al., 2017).

**Figure 1:**
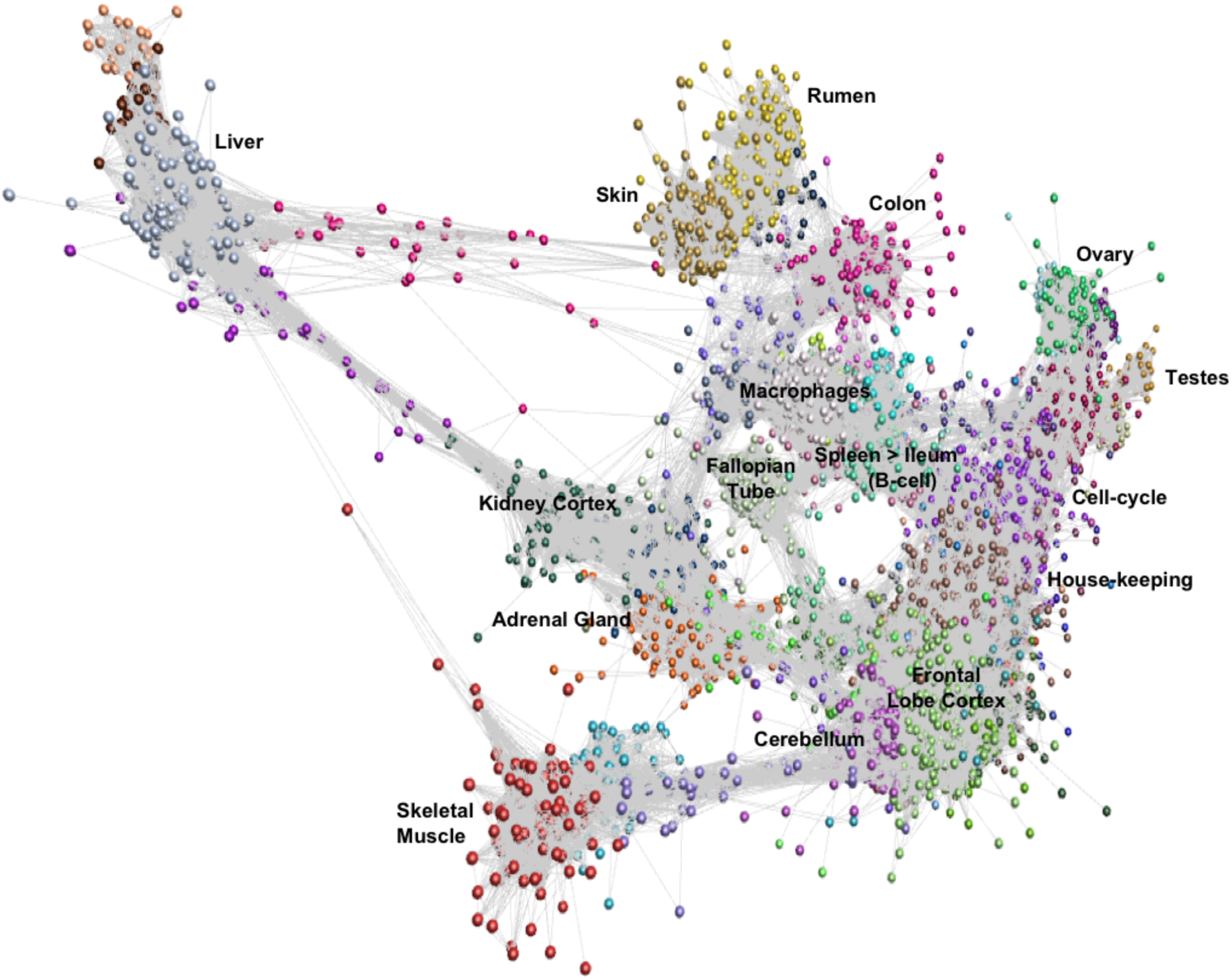
Gene-to-gene network graph of the goat mini-atlas dataset. Each ‘node’ represents a gene and each ‘edge’ represents correlations between individual measurements above the set threshold. The graph comprised 16,172 nodes (genes) and 1,574,259 edges (Pearson correlations ≥ 0.83), MCL inflation = 2.2, Pearson Product Correlation Co-efficient = 0.83.

**Table 3:** Annotation of the 20 largest network clusters in the goat mini-atlas dataset (> indicates decreasing expression profile).

| Cluster ID | Number of Genes | Profile Description | Class | Enriched GO terms |
| --- | --- | --- | --- | --- |
| 1 | 1795 | Cortex>cerebellum | Brain | cognition, neurotransmitter transport, synaptic transmission |
| 2 | 1395 | Thymus>Spleen>Ileum | Cell-Cycle | DNA-dependent DNA replication, DNA repair |
| 3 | 795 | General | House Keeping | mRNA processing, regulation of RNA splicing |
| 4 | 505 | Liver | Oxidative-Phosphorylation | oxidation-reduction process, fatty acid oxidation |
| 5 | 494 | General | House Keeping | RNA binding, nucleolus |
| 6 | 481 | Testes | Male Reproduction | male meiosis, spermatogenesis |
| 7 | 449 | Skin > Rumen | Epithelial | skin morphogenesis, keratinocyte differentiation |
| 8 | 374 | Fallopian Tube | Motile Cilia | motile cilium, ciliary basal body |
| 9 | 351 | Skeletal muscle | Muscle | muscle fibre development, motor activity |
| 10 | 337 | Spleen>Ileum | Immune | immune response, B-cell activation, cytokine activity |
| 11 | 290 | Macrophages | Immune | response to lipopolysaccharide, phagocytic vesicle |
| 12 | 241 | Colon Large | Gastrointestinal tract | microvillus, actin filament bundle |
| 13 | 226 | Rumen > Skin | Gastrointestinal/Epithelial | epidermis development, chloride channel activity |
| 14 | 219 | Adrenal Gland | Endocrine | oxidation-reduction process, sterol metabolic process |
| 15 | 211 | BMDMs | Fibroblasts | collagen binding, positive regulation of fibroblast proliferation |
| 16 | 134 | General | Ribosomal | ribosomal large subunit biogenesis, ribosome |
| 17 | 133 | Kidney Cortex | Mesoendonephric organogenesis | sodium ion homeostasis, skeletal system morphogenesis |
| 18 | 119 | Ovary | Oogenesis | growth factor activity, nucleosome disassembly |
| 19 | 113 | Ileum>Spleen>Thymus | Immune | B-cell proliferation, cytokine activity |
| 20 | 108 | Uterus, Uterine Horn | Organogenesis | tissue remodelling, bone morphogenesis |

The largest of the clusters (Cluster 1) contained 1795 genes that were almost exclusively expressed in the central nervous system (cortex, cerebellum) reflecting the high transcriptional activity and complexity in the brain. Significant GO terms for cluster 1 included cognition (p=4.6×^-17^) and synaptic transmission (p=2.5×10^-30^). Other tissue-specific clusters; e.g. 4 (liver), 6 (testes), 7 (skin/rumen), 14 (adrenal) and 17 (kidney) were similarly enriched for genes associated with known tissue-specific functions. In each case, the likely function of unannotated protein-coding genes within these clusters could be inferred by association with genes of known function that share the same cell or tissue specific expression pattern. Cluster 9 showed a high level of tissue specificity and included genes associated with skeletal muscle function and development including MSTN which encodes a protein that negatively regulates skeletal muscle cell proliferation and differentiation (Wang et al., 2012). Several myosin light and heavy chain genes (e.g. MYH1 and MYL1) and transcription factors that are specific to muscle including (MYOG and MYOD1) were also found in cluster 9. GO terms for muscle were enriched in cluster 9 e.g. muscle fiber development (p=3.8×10^-13^) and structural constituent of muscle (p=1.8×10^-11^). Genes expressed in muscle are of particular biological and commercial interest for livestock production and represent potential targets for gene editing (Yu et al., 2016). Cluster 8 was also highly tissue specific and included genes expressed in the fallopian tube with enriched GO terms for cilium movement (p=1.4×10^-15^) and cilium organization (p=2.3×10^-15^). A motile cilia cluster was identified in the fallopian tube in the sheep gene expression atlas (Clark et al., 2017) and a similar cluster was enriched in chicken in the trachea (Bush et al., 2018a). The goat mini-atlas also included several clusters that were enriched for immune tissues and cell types and we have based our analysis in part upon the premise that the greatest differences between small ruminant species likely involve the immune system.

### Gene expression in the neonatal gastrointestinal tract

Three regions of the gastrointestinal (GI) tract were sampled; the ileum, colon and rumen. These regions formed distinct clusters in the network graph. The genes comprising these clusters were highly correlated with the physiology of the tissues. Goats are ruminant mammals and at one-week of age (when tissues were collected) the rumen is vestigial. Even at this early stage of development, the typical epithelial signature of the rumen (Xiang et al., 2016a; Xiang et al., 2016b) was observed. Genes co-expressed in the rumen (clusters 7 and 13 – Table 3) were typical of a developing rumen epithelial signature (Bush et al., 2019) and were associated with GO terms for epidermis development (p=0.00016), keratinocyte differentiation (p=1.5×10^-14^) and skin morphogenesis (p=8.2×10^-6^). Large colon (cluster 12) included several genes associated with GO terms for microvillus organization (p=1×10^6^) and microvillus (p=6.3×10^6^) including MYO7B which is found in the brush border cells of epithelial microvilli in the large intestine. The microvilli function as the primary surface of nutrient absorption in the gastrointestinal tract, and as such numerous phospholipid-transporting ATPases and solute carrier genes were found in the large colon cluster.

Throughout the GI tract there was a strong immune signature, similar to that observed in neonatal and adult sheep (Bush et al., 2019), which was greatest in clusters 10 and 19 (Table 3) where expression was high in the ileum and Peyer’s patches, thymus and spleen. Cluster 10 had a more general immune related profile with higher expression in the spleen and significant GO terms associated with cytokine receptor activity (p=1.3×10^-8^) and T cell receptor complex (p=0.00895). Several genes involved in the immune and inflammatory response were found in cluster 10 including CD74, IL10 and TLR10. The expression pattern for cluster 19 was associated with B-cells including GO terms for B cell proliferation (p=1.4×10^-7^), positive regulation of B cell activation (p=4.9×10^-6^) and cytokine activity (p=0.0051). Genes associated with the B-cell receptor complex CD22, CD79B, CD180 and CR2, and interleukins IL21R and IL26 were expressed in cluster 19 (Treanor, 2012). This reflects the fact that we sampled the Peyer’s patch with the ileum, which is a primary lymphoid organ of B-cell development in ruminants (Masahiro et al., 2006).

Each of the GI tract clusters included genes associated with more than one cell type/cellular process. This complexity is a consequence of gene expression patterns from the lamina propria, one of the three layers of the mucosa. The lamina propria lies beneath the epithelium along the majority of the GI tract and comprises numerous different cell types from endothelial, immune and connective tissues (Ikemizu et al., 1994). This gene expression pattern, which is also observed in sheep (Clark et al., 2017; Bush et al., 2019) and pigs (Freeman et al., 2012), highlights the complex multi-dimensional physiology of the ruminant GI tract.

### Macrophage-associated signatures

A strong immune response is vitally important to neonatal mammals. Macrophages constitute a major component of the innate immune system acting as the first line of defense against invading pathogens and coordinating the immune response by triggering anti-microbial responses and other mediators of the inflammatory response (Hume, 2015). Several clusters in the goat mini-atlas exhibited a macrophage-associated signature. Cluster 11 (Table 3), contained several macrophage marker genes, including CD68 which is expressed in AMs and BMDMs. The cluster includes the macrophage growth factor, CSF1, indicating that as in sheep (Clark et al., 2017), pigs (Freeman et al., 2012) and humans (Schroder et al., 2012) but in contrast to mice, goat macrophages are autocrine for their own growth factor. GO terms associated with cluster 11 included phagocytosis (p=3.5×10^-10^), inflammatory response (p=1.4×10^-8^) and cytokine receptor activity (p=0.00031). Many of the genes that were up-regulated in AMs in cluster 11, including C-type lectins CLEC4A and CLEC5A, have been shown to be down regulated in sheep (Clark et al., 2017; Bush et al., 2019), pigs (Freeman et al., 2012) and humans (Baillie et al., 2017) in the wall of the intestine. This highlights functional transcriptional differences in macrophage populations. AMs respond to microbial challenge as the first line of defense against inhaled pathogens. In contrast, macrophages in the intestinal mucosa down-regulate their response to microorganisms as a continuous inflammatory response to commensal microbes would be undesirable.

Cluster 11 (Table 3) also included numerous pro-inflammatory cytokines and chemokines which were up-regulated following challenge with lipopolysaccharide (LPS). Response to LPS was also reflected in several significant GO terms associated with this cluster including, cellular response to lipopolysaccharide (p=5.8×10^-10^) and cellular response to cytokine stimulus (p=9.5×10^-8^). C-type lectin CLEC4E, which is known to be involved in the inflammatory response (Baillie et al., 2017), interleukin genes such as IL1B and IL27, and ADGRE1 were all highly inducible by LPS in BMDMs. ADGRE1 (EMR1,F4/80) is a monocyte-macrophage marker involved in pattern recognition which exhibits inter-species variation both in expression level and response to LPS stimulation (Waddell et al., 2018). Based upon RNA-Seq data, ruminant genomes were found to encode a much larger form of ADGRE1 than monogastric species, with complete duplication of the extracellular domain [44].

### Comparative analysis of macrophage-associated transcriptional responses in sheep and goats

Transcriptional differences are linked to species-specific variation in response to disease, and have been widely documented in livestock (Bishop and Woolliams, 2014). For instance, ruminants differ in their response to a wide range of economically important pathogens. Variation in the expression of NRAMP1 (SLC11A1) is involved in the response of sheep and goat to Johne’s disease (Cecchi et al., 2017). Similarly, resistance to *Haemonchus contortus* infections in sheep and goats is associated with a stronger Th2-type transcriptional immune response (Gill et al., 2000; Alba-Hurtado and Munoz-Guzman, 2013). To determine whether goats and sheep differ significantly in immune transcriptional signatures we performed a comparative analysis of the macrophage samples from the goat mini-atlas and those included in our gene expression atlas for sheep (Clark et al., 2017). One caveat to this analysis that should be noted is that the sheep and goat samples were unfortunately not age-matched and as such differences in gene expression could be an effect of developmental stage rather than species-specific differences. However, as macrophage samples from both species were kept in culture prior to collection and analysis we would expect the effect of developmental stage to be minimal.

We performed differential analysis of genes expressed in goat and sheep AMs (Supplementary Table S13). The top 25 genes up- and down-regulated in goat relative to sheep based on log2FC are shown in Fig 2. Several genes involved in the inflammatory and immune response including, interleukins IL33 and IL1B and C-type lectin CLEC5A were up-regulated in goat AMs relative to sheep. In contrast those that were down regulated in goat relative to sheep did not have an immune function but were associated with more general physiological processes. This may reflect species-specific differences but could also indicate that the immune response in AMs is age-dependent i.e. neonatal animals exhibit a primed immune response while a more subdued response is exhibited by adult sheep whose adaptive immunity has reached full development.

**Figure 2:**
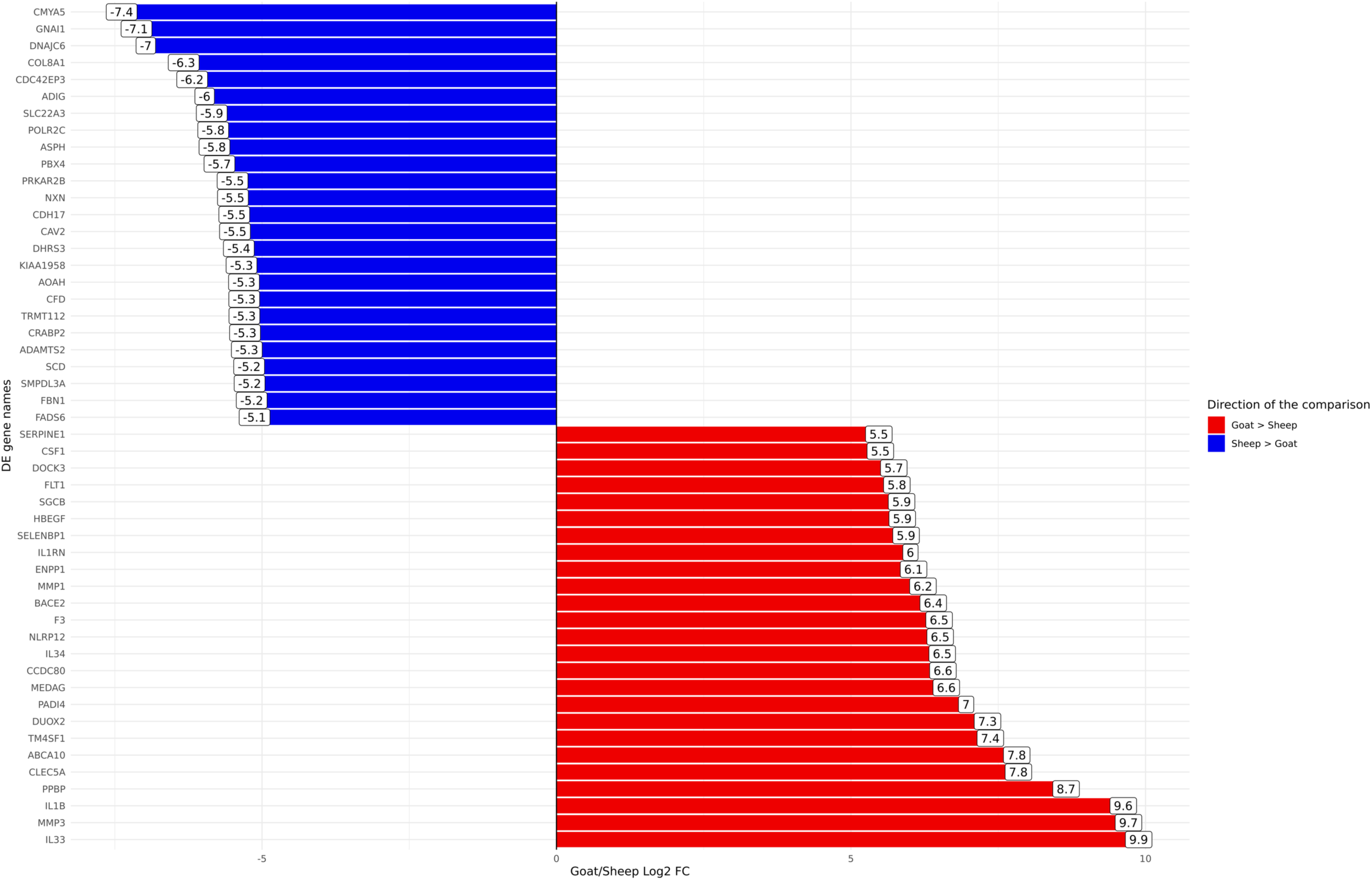
Differentially expressed genes (FDR<10%) between goat and sheep alveolar macrophages. The top 25 up-regulated in goat relative to sheep (red) and the top 25 down-regulated in goat relative to sheep (blue) are shown.

Using differential expression analysis (Robinson et al., 2010) we also compared the gene expression estimates for sheep and goat BMDMs (+/-) LPS, to compile a list of genes for each species that were up or down regulated in response to LPS (Supplementary Table S14A goat and Supplementary Table S14B sheep). These lists were then merged using the methodology described above (see Methods section) to highlight genes that differed in their response to LPS between the two species. In total 188 genes exhibited significant differences between goats and sheep (FDR<10%, Log2FC>=2) in response to LPS (Supplementary Table S15). The genes which showed the highest level of dissimilarity in response to LPS between goats and sheep (Dis_Index>=2) are illustrated in Fig 3. Several immune genes were upregulated in both goat and sheep BMDMs in response to LPS stimulation but differed in their level of induction between the two species (top right quadrant Fig 3). IL33, IL36B, PTX3, CCL20, CSF3 and CSF2 for example, exhibited higher levels of induction in sheep BMDMs relative to goat, and vice versa for ICAM1, IL23A, IFIT2, TNFSF10, and TNFRSF9. Several genes were upregulated in sheep but downregulated in goat BMDMs (e.g. KIT) (top left quadrant Fig 3), and upregulated in goat, but downregulated in sheep (e.g. IGFBP4) (bottom right quadrant Fig 3).

**Figure 3:**
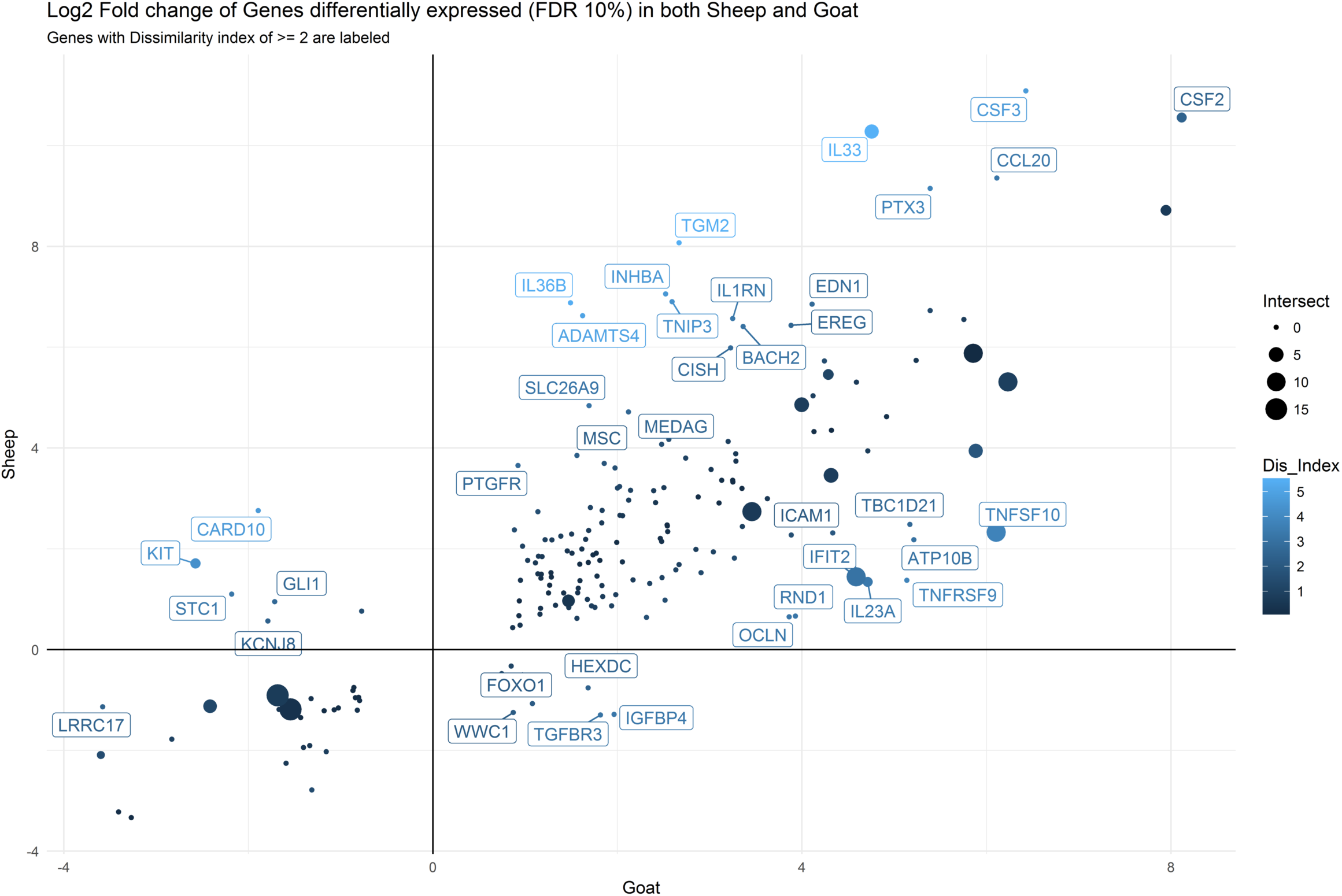
Comparative analysis of differentially expressed genes (FDR<10%, Log2FC>=2) in goat and sheep BMDM. The genes which showed the highest level of dissimilarity in response to LPS between goats and sheep (Dis_Index>=2) are shown. Top right quadrant: genes that were up-regulated in both goat and sheep but differed in their level of induction between the two species. Top left quadrant: genes that were up-regulated in sheep but down-regulated in goat. Bottom right quadrant: genes up-regulated in goat, but down-regulated in sheep.

Overall the transcriptional patterns in BMDMs stimulated with LPS were broadly similar between the two species. Some interesting differences in individual genes were observed that could contribute to species-specific responses to infection. For instance, IL33 and IL23A both exhibited a higher level of induction in sheep BMDMs after stimulation with LPS relative to goat (Fig 3). In humans IL33 has a protective role in inflammatory bowel disease by inducing a Th2 immune response (Lopetuso et al., 2013). An enhanced Th2 response, which accelerates parasite expulsion, has been associated with *H. contortus* resistance in sheep (Alba-Hurtado and Munoz-Guzman, 2013). Conversely, higher expression of IL23A is associated with susceptibility to *Teladorsagia circumcincta* infection (Gossner et al., 2012). Little is known about the function of IL33 and IL23A in goats. They are members of the interleukin-1 family which play a central role in the regulation of immune and inflammatory response to infection (Dinarello, 2018). Given the similarities in their expression patterns, it is reasonable to assume that these genes are regulated in a similar manner to sheep and involved in similar biological pathways. As such they would be suitable candidate genes to investigate further to determine if they underlie species-specific variation in susceptibility to pathogens (Bishop and Stear, 2003; Bishop and Morris, 2007).

### Expression patterns of genes associated with functional traits in goats

The goat mini-atlas dataset is a valuable resource that can be used by the livestock genomics community to examine the expression patterns of genes of interest that are relevant to ruminant physiology, immunity, welfare, production and adaptation/resilience particularly in tropical agri-systems. Several genes, associated with functional traits in goats, have been identified using genome wide association studies (GWAS). Insulin-like growth factor 2 (IGF2), for example, is associated with growth rate in goats (Burren et al., 2016), and was highly expressed in tissues with a metabolic function including, kidney cortex, liver and adrenal gland (Fig 4A). As expected expression of myostatin (MSTN), which encodes a negative regulator of skeletal muscle mass, was highest in skeletal muscle in comparison with the other tissues (Fig 4B). MSTN is a target for gene-editing in goats to promote muscle growth (e.g. Yu et al., 2016). Expression of genes associated with fecundity and litter size in goats, including GDF9 and BMPR1B (Feng et al.; Shokrollahi and Morammazi, 2018), were highest in the ovary (Fig 4C & D). The ovary included here is from a neonatal goat and these results correlate with similar observations in sheep where genes essential for ovarian follicular growth and involved in ovulation rate regulation and fecundity were highly expressed in foetal ovary at 100 days gestation (Clark et al., 2017).

**Figure 4:**
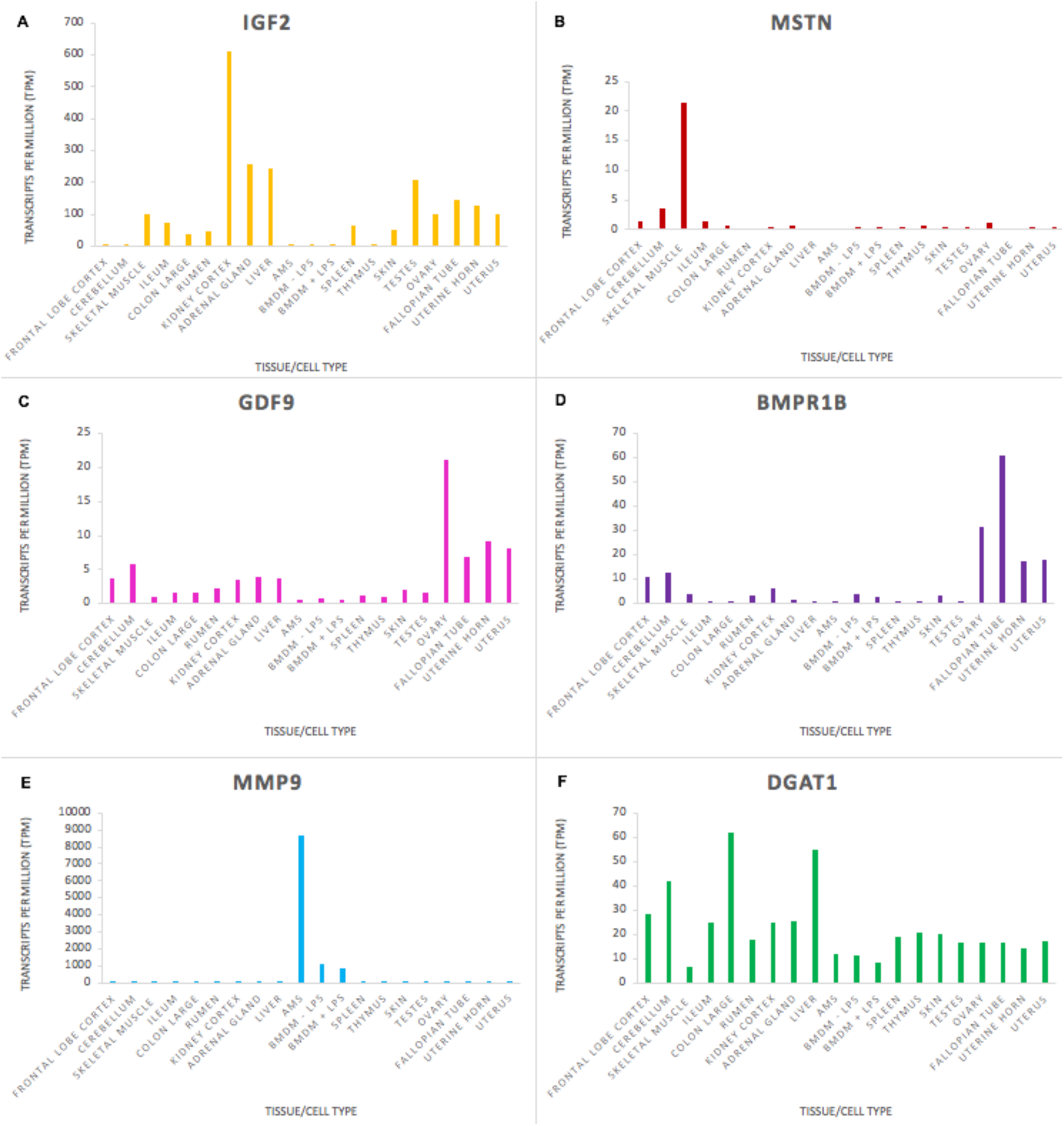
Expression levels (transcripts per million) of genes involved in functional traits in goats to illustrate tissue and cell type or ubiquitous expression patterns. A: IGF2 is associated with growth rate; B: MSTN is associated with muscle characteristics; C: GDF9 is associated with ovulation rate; D: BMPR1 is associated with fecundity; E: MMP9 is associated with resistance to mastitis; F: DGAT1 is associated with fat content in goat milk.

Some genes, particularly those involved in the immune response had high tissue or cell type specific expression. Matrix metalloproteinase-9 (MMP9), which is involved in the inflammatory response and linked to mastitis regulation in goats (Li et al., 2016) was very highly expressed in macrophages, particularly AMs, in comparison with other tissues (Fig 4E). Other genes that are important for goat functional traits were fairly ubiquitously expressed. The expression level of Diacylglycerol O-Acyltransferase 1 (DGAT1) which is associated with milk fat content in dairy goats (Martin et al., 2017) did not vary hugely across the tissues sampled (Fig 4F), although there was slightly higher expression in some tissues (e.g. colon and liver) relative to immune tissues (e.g. thymus and spleen). DGAT1 encodes a key metabolic enzyme that catalyses the last, and rate-limiting step of triglyceride synthesis, the transformation from a diacylglycerol to a triacylglycerol (Bell and Coleman, 1980). This is an important cellular process undertaken by the majority of cells, explaining its ubiquitous expression pattern. Two exonic mutations in the DGAT1 gene in dairy goats have been associated with a notable decrease in milk fat content (Martin et al., 2017). Understanding how these, and other variants for functional traits, are expressed can help us to determine how their effect on gene expression and regulation influences the observed phenotypes in goat breeding programmes.

### Allele-specific expression

Using mapping bias correction for robust positive ASE discovery (Salavati et al., 2019), we were able to profile moderate to extreme allelic imbalance across tissues and cell types, at the gene level, in goats. The raw ASE values for every tissue/cell type are included in Supplementary Dataset S4. We first calculated the distribution of heterozygote sites per gene, as a measure of homogeneity of input sites, and found there was no significant difference between the eight individual goats included in the study (Supplementary Fig S2).

Several genes exhibited pervasive allelic imbalance (i.e. where the same imbalance in expression is shared across several tissues/cell types) (Fig 5). For example, allelic imbalance was observed in the mitochondrial ribosomal protein MRPL17 in 16 tissues/cell types (except skeletal muscle and rumen). SERPINH1, a member of the serpin superfamily, was the only gene in which an imbalance in expression was detected in all tissues/cell types. Allelic imbalance was observed in COL4A1 in 11 tissues, and was highest in the rumen and skin samples. COL4A1 has been shown to be involved in the growth and development of the rumen papillae in cattle (Nishihara et al., 2018) and sheep (Bush et al., 2019). The highest levels of allelic imbalance in individual genes were observed in ribosomal protein RPL10A in ileum and SPARC in liver (Fig 5).

**Figure 5:**
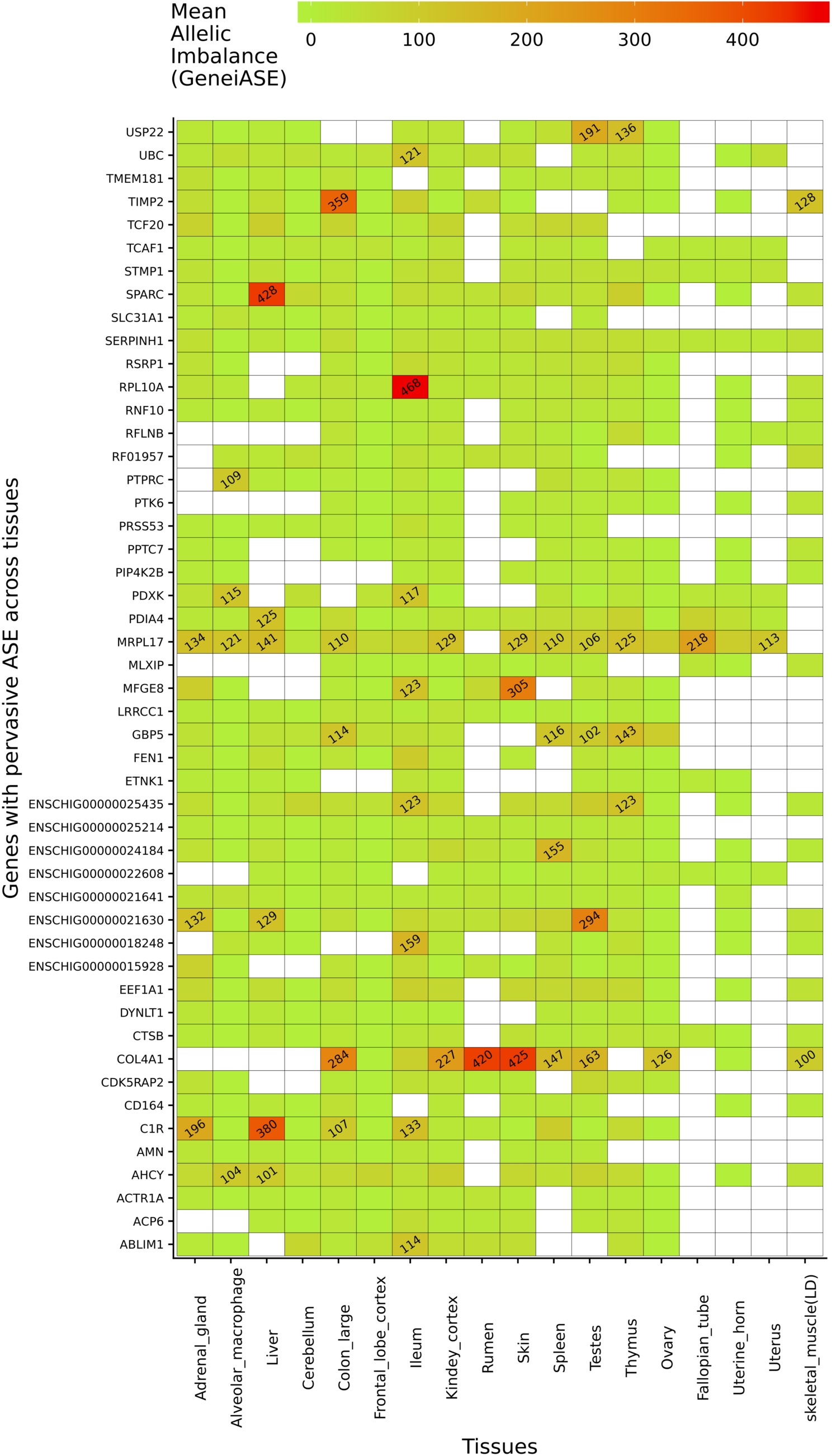
Genes exhibiting the largest mean allelic imbalance. (i.e. allele-specific expression averaged across all heterozygote sites within each gene) across 17 tissues and one cell type from the goat mini-atlas dataset visualised as a heatmap (red indicating the highest level of mean allelic imbalance and green the least).

The ASE profiles were highly tissue- or cell type-specific, with strong correlations between samples from the same organ system (Fig 6). For example, ASE profiles in female reproductive system (ovary, fallopian tube, uterine horn, uterus), GI tract (colon and ileum) and brain (cerebellum and frontal lobe cortex) tissues were highly correlated. The two tissues showing the largest proportion of shared allele-specific expression were the ovary and liver (Fig 6). This might reflect transcriptional activity in these tissues in neonatal goats during oogenesis (ovary) and haematopoiesis (liver). Future work could determine if these ASE patterns were observed at other stages of development, or whether they are time-dependant.

**Figure 6:**
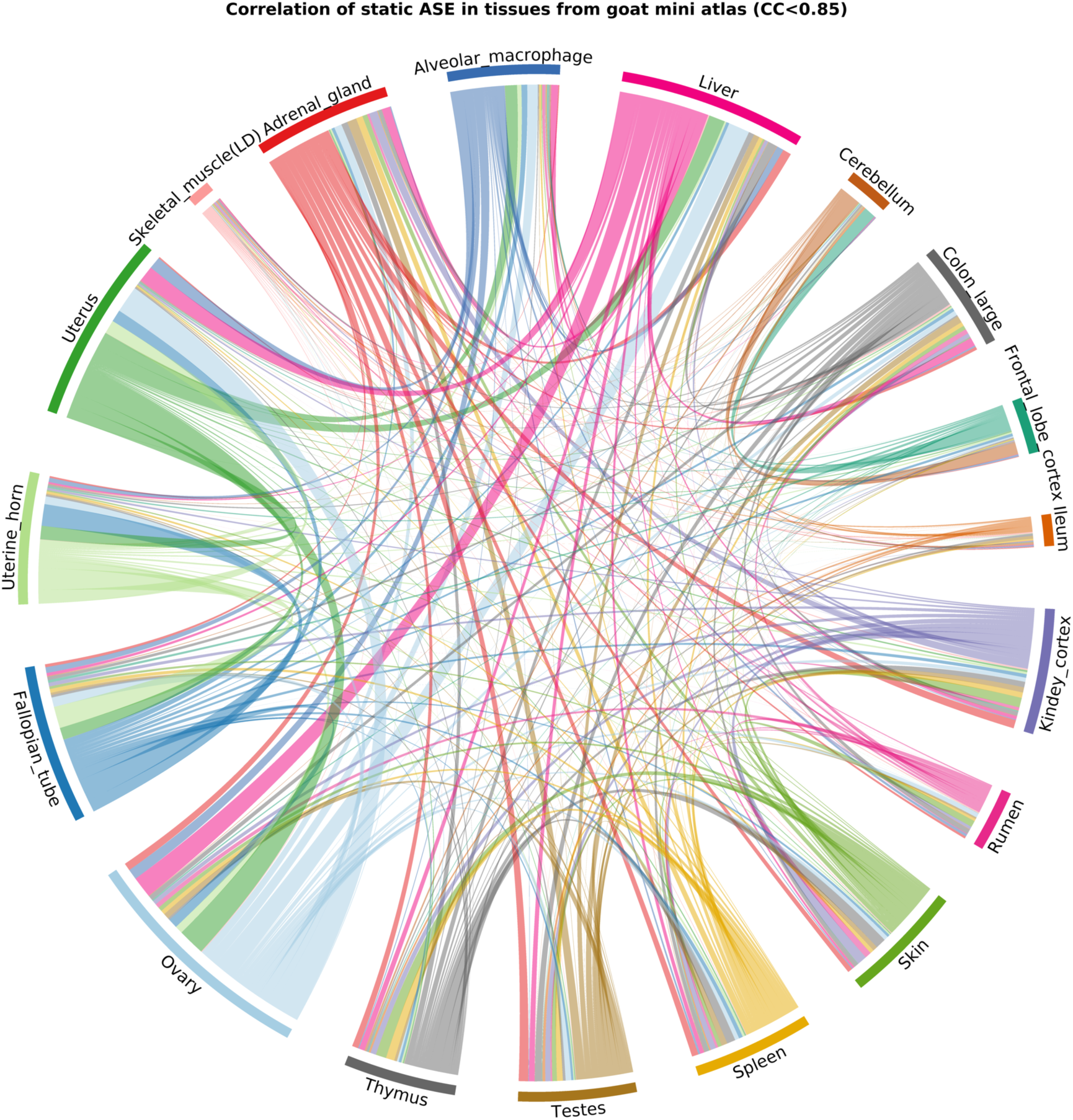
Correlation of ASE profiles shared across tissues/cell types from the goat mini-atlas dataset. Each section represents the genes showing significant allelic imbalance within the tissue. The chords represent the correlation coefficient (CC<0.85) of ASE profiles shared between the samples (i.e. the proportion of genes showing co-imbalance).

The next step of this analysis would be to analyse ASE at the variant (SNV) level. This would allow us to identify variants driving ASE and determine whether they were located within important genes for functional traits. These variants could then be weighted in genomic prediction algorithms for genomic selection, for example. The sequencing depth used for the goat mini-atlas is, however, insufficient for statistically robust analysis at the SNV level. Nevertheless, it does provide a foundation for further analysis of ASE relevant to functional traits using a suitable dataset, ideally from a larger number of individuals (e.g. for aseQTL analysis (Wang et al., 2018)) and at a greater depth.

## Conclusions

We have created a mini-atlas of gene expression for the domestic goat. This expression dataset complements the genetic and genomic resources already available for goat (Tosser-Klopp et al., 2014; Stella et al., 2018; Talenti et al., 2018), and provides a set of functional information to annotate the current reference genome (Bickhart et al., 2017; Worley, 2017). We were able to detect the majority (90%) of the transcriptome from a subset of 22 transcriptionally rich tissues and cell-types representing all the major organ systems, providing proof of concept that this mini-atlas approach is useful for studying gene expression and for functional annotation. Using the mini-atlas dataset we annotated 15% of the unannotated genes in ARS1. Our dataset was also used by the Ensembl team to create a new gene build for the goat ARS1 reference genome (https://www.ensembl.org/Capra_hircus/Info/Index).

We have also provided transcriptional profiling of macrophages in goats and a comparative analysis with sheep. This provides a foundation for further analysis in more tissues and cell types in age-matched animals, and in disease challenge experiments for example. Prior to this study little was known about the transcription in goat macrophages. While more information is available on goat monocyte derived macrophages (Adeyemo et al., 1997; Taka et al., 2013; Walia et al., 2015), there was previously relatively little knowledge available on the characteristics of goat BMDMs. In addition, few reagents are available for immunological studies in goat, with most studies relying on cross-reactivity with sheep and cattle antibodies (Entrican, 2002; Hope et al., 2012). Recently a characterisation of goat antibody loci has been published using the new reference genome ARS1 (Schwartz et al., 2018), demonstrating the usefulness of a highly contiguous reference genome with high quality functional annotation for the development of new resources for livestock species. The goat mini-gene expression atlas complements the large gene expression dataset we have generated for sheep and contributes to the genomic resources we are developing for interpretation of the relationship between genotype and phenotype in small ruminants.

## Data Availability

We have made the files containing the expression estimates for the goat mini-atlas (Supplementary Dataset S2 (unaveraged) and Supplementary Dataset S3 (averaged)) available for download through the University of Edinburgh DataShare portal (https://doi.org/10.7488/ds/2591). Sample metadata for all the tissue and cell samples collected has been deposited in the EBI BioSamples database under project identifier GSB-2131 (https://www.ebi.ac.uk/biosamples/samples/SAMEG330351) according to FAANG metadata and data sharing standards. The raw fastq files for the RNA-Seq libraries are deposited in the European Nucleotide Archive (https://www.ebi.ac.uk/ena) under the accession number PRJEB23196. The data submission to the ENA includes experimental metadata prepared according to the FAANG Consortium metadata and data sharing standards. The BAM files are also available as analysis files under accession number PRJEB23196 (‘BAM file 1’ are mapped to the NCBI version of ARS1 and ‘BAM file 2’ to the Ensembl version). The data from sheep included in this analysis has been published previously and is available via (Clark et al., 2017) and under ENA accession number PRJEB19199. Details of all the samples for both goat and sheep are available via the FAANG data portal (http://data.faang.org/home). All experimental protocols are available on the FAANG consortium website at http://www.ftp.faang.ebi.ac.uk/ftp/protocols

## Author Contributions

ELC, CM and DAH designed the study. MA, AD and DAH provided guidance on project design, sample collection and analysis. DAH, MA and AD secured the funding for the project with CM. CM and ELC collected the samples with ZL and MEBM who performed the post mortems. CM performed the RNA extractions. SJB performed the bioinformatic analyses. MS performed the analysis of allele-specific expression and assisted CM with the comparative analysis. CM performed the network cluster analysis with ELC. CM and ELC wrote the manuscript. All authors contributed to editing and approved the final version of the manuscript.

## Acknowledgements

The authors would like to thank Lindsey Waddell, Anna Raper, Rahki Harne, Rachel Young, Lucas Lefevre and Lucy Freem for assistance with isolating and characterising BMDMs. Peter Harrison and Jun Fan at the FAANG Data Coordination Centre provided advice on upload of raw data, sample and experimental metadata to the ENA and BioSamples.

## Ethics approval and consent to participate

Approval was obtained from The Roslin Institute, University of Edinburgh’s Animal Work and Ethics Review Board (AWERB). All animal work was carried out under the regulations of the Animals (Scientific Procedures) Act 1986.

## Supplemental Figures

S1 Figure: Sample-to-sample network graph of the samples included in the goat mini-atlas dataset.

S2 Figure: Distribution of heterozygote (bi-allelic) sites per genes for each of the eight individual goats included in the study. The bi-allelic sites were compared to the ARS1 Ensembl v96 reference variant call format (VCF) track which includes 22,379 genes and more than 12 million heterozygote sites. On average for each animal 3,004,867 heterozygote loci were examined for allelic imbalance. The ARS1 reference (Ref) distribution is shown in blue with the distribution for each individual goat included in this study (male m1-7, female f8) overlaid in red.

### Supplemental Datasets

S1 Dataset: Gene expression estimates for AMs and BMDMs (+/- LPS) unaveraged across biological replicates for the subset of sheep gene expression atlas dataset included for comparative analysis.

S2 Dataset: Gene expression estimates unaveraged across biological replicates for the goat mini-atlas dataset.

S3 Dataset: Gene expression estimates averaged across biological replicates for the goat mini-atlas dataset.

S4 Dataset: Estimates of allele-specific expression for each sample from the goat mini-atlas dataset using the GeneiASE model.

### Supplemental Tables

S1 Table: Quantity and quality measurements of isolated RNA from all tissue and cell-types in the goat mini-atlas dataset.

S2 Table: Summary of transcript models generated using the HISAT2/stringtie pipeline in comparison with gene models in the reference genome ARS1.

S3 Table: Novel transcript models generated for goat using the HISAT2/stringtie pipeline. S4 Table: A list of all undetected genes in the goat mini-atlas dataset.

S5 Table: A list of all undetected transcripts in the goat mini-atlas dataset.

S6 Table: The proportion of transcripts with detectable expression (TPM >1) in the goat mini-atlas relative to the ARS1 reference transcriptome at the gene level.

S7 Table: The proportion of transcripts with detectable expression (TPM >1) in the goat mini-atlas relative to the ARS1 reference transcriptome at the transcript level.

S8 Table: A short-list containing a conservative set of gene annotations using the goat mini-atlas dataset.

S9 Table: The ‘long’ list of genes annotated using the goat mini-atlas dataset.

S10 Table: A list of unannotated genes associated with a gene description, but not necessarily an HGNC symbol.

S11 Table: Genes included in each cluster from the network cluster analysis of the goat mini-atlas dataset.

S12 Table: GO term enrichment of each of the clusters from the network cluster analysis of the goat mini-atlas dataset.

S13 Table: Differentially expressed genes in goat and sheep alveolar macrophages.

S14 Table: Differentially expressed genes in goat (A) and sheep (B) bone marrow derived macrophages (BMDM) (+/-) LPS.

S15 Table: Genes that exhibited significant differences between goats and sheep (FDR<10%, Log2FC>=2) in response to L

