## Supplementary figures and images for "A mini-atlas of gene expression for the domestic goat (*Capra hircus*) reveals transcriptional differences in immune signatures between sheep and goats"

### Supplementary Figure S1

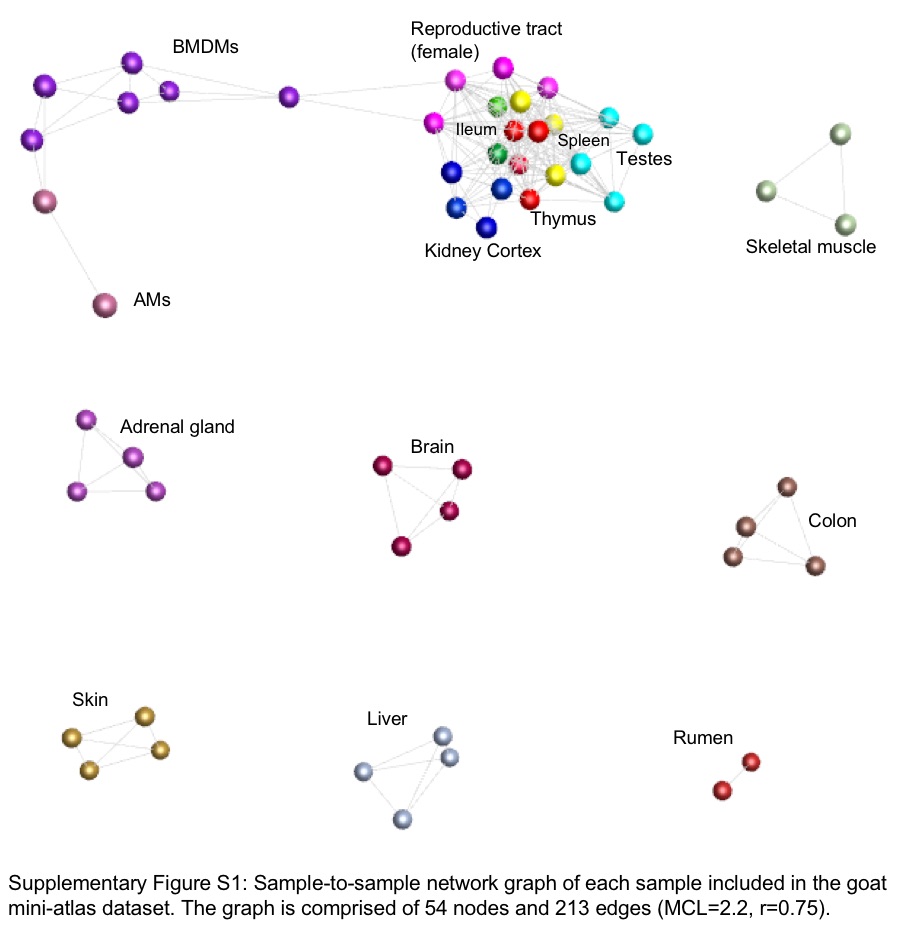

### Supplementary Figure S2

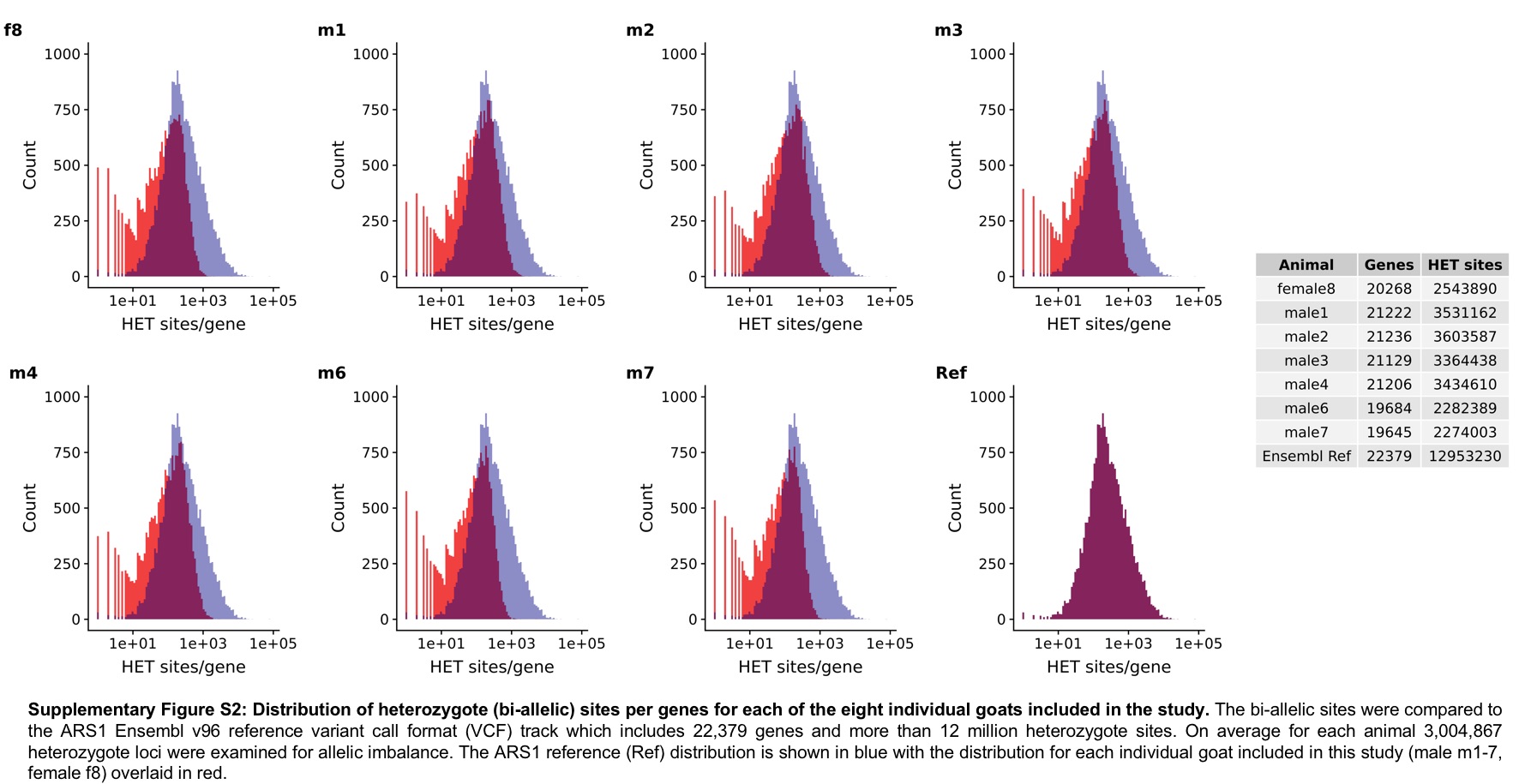
